# Blind cavefish evolved food-searching behavior without changing sensory modality compared with sighted conspecies in the dark

**DOI:** 10.1101/2023.06.12.544672

**Authors:** Kyleigh Kuball, Vânia Filipa Lima Fernandes, Daisuke Takagi, Masato Yoshizawa

## Abstract

In nature, animals must navigate to forage according to their sensory inputs. Different species use different sensory modalities to locate food efficiently. For teleosts, food emits visual, mechanical, chemical, and/or possibly weak-electrical signals, which can be detected by optic, auditory/lateral line, and olfactory/taste buds sensory systems. However, how fish respond to and use different sensory inputs when locating food, as well as the evolution of these sensory modalities, remain unclear. We examined the Mexican tetra, *Astyanax mexicanus*, which is composed of two different morphs: a sighted riverine (surface fish) and a blind cave morph (cavefish). Compared with surface fish, cavefish have enhanced non-visual sensory systems, including the mechanosensory lateral line system, chemical sensors comprising the olfactory system and taste buds, and the auditory system to help navigate toward food sources. We tested how visual, chemical, and mechanical stimuli evoke food-seeking behavior. In contrast to our expectations, both surface fish and cavefish did not follow a gradient of chemical stimulus (food extract) but used it as a cue for the ambient existence of food. Surface fish followed visual cues (red plastic beads and food pellets), but, in the dark, were likely to rely on mechanosensors—the lateral line and/or tactile sensor—as cavefish did. Our results indicate cavefish used similar sensory modality to surface fish in the dark, while adherence levels to stimuli were higher in cavefish. In addition, cavefish evolved an extended circling strategy to capture food, which may yield a higher chance to capture food by swimming-by the food multiple times instead of once through zigzag motion. In summary, we propose ancestors of cavefish similar to surface fish may have needed little modification in food-seeking strategy to adapt to the dark.

## Introduction

Many teleost species rely on visual information for foraging, although fishes employ a wide range of sensory modalities for foraging strategies [1–4]. These strategies range from drift-hunting by coelacanths that use a single sensory modality (electroreception) to detect benthic prey [5], to the multi-sensory, active pursuit of prey by bonnethead sharks, which use long-distance olfactory signals followed by visual cues to precisely locate prey [2].

Given the breadth of sensory systems, how the coordination and hierarchical use of sensory systems change during the adaptation to a new environment remains unclear. Depending on species, different mechanisms are favored, such as mechano-, chemo-, and/or electro-sensing [1,2]. For foraging tradeoffs between finding (energy loss) and consuming food gains (energy gain), animals should strategize to maximize energy gain with minimum loss by leveraging available sensory inputs [6].To tackle this question, we chose the freshwater Mexican tetra, *Astyanax mexicanus. Astyanax mexicanus* is a ∼6 cm freshwater fish, consisting of two morphs: riverine and sighted surface form (surface fish: colonizing in a rage of south Texas USA to the south American continent) and the cave-dwelling blind form (cavefish: limestone mountain ranges at Northeast Mexico). We then conducted foraging experiments comparing these different populations of the same species.

Cavefish show higher responses to mechanical vibration stimulus at ∼40 Hz than surface fish. The 40 Hz vibration can be typically generated by crawling crustaceans [7] which is promoted by the increased cranial mechanosensory lateral line. Fish with higher vibration responses, called vibration attraction behavior (VAB), dominated over prey capture in the dark [8,9]. Cavefish also have finer chemical sensing, such as the ability to respond to 10^5^ lower concentrations of amino acids than surface fish (i.e., cavefish can respond to 10^−10^ M of alanine, whereas surface fish respond to 10^−5^ M of it or higher) [10]. In contrast, no detectable difference in auditory response has been reported between surface fish and cavefish [11] and there is no comparative study in tactile sensing between these two morphs (but see Voneida & Fish [12]).

Upon this powerful comparative model system, it remains largely unknown how these sensory systems were strategically utilized during foraging: are these sensory systems used equally for foraging, or is there any hierarchical order of the usage of the sensory systems? Then, if there is a hierarchical order, what is its ecological relevance? To provide answers to these questions, we designed experiments using varying stimuli. We used (1) water droplets as the source of mechanical stimulus (auditory only, when it hits the water surface), (2) food extract suspended in water as the source of the mechanical (auditory) + chemical stimuli—only chemical stimulus is the additional to (1), (3) red plastic beads as visual + mechanical (auditory + lateral line/tactile) stimuli, which are additional to (1), (4) food extract and plastic beads, and (5) fish commercial diet as a positive control. We then measured latency as the initial response to these stimuli, number of foraging attempts as the proxy for robustness of foraging mode, and zigzag and circling measurements (duration and bout numbers) to characterize two foraging strategies in surface fish and cavefish. Foraging with circling is typical in cavefish; however, it was not clear if surface fish showed zigzag or circling in the dark before this study (see Result and Discussion section about the behavioral characteristics of zigzag and circling).

Our result indicated that, for latency measurements, surface fish did not respond to sole auditory stimulus (water droplet) in either light or dark conditions, but cavefish did, suggesting surface fish require multiple sensory inputs. In contrast, the cavefish foraging behavior could be driven by auditory stimulus alone. Object stimuli (beads) evoked slightly higher foraging behavior in both surface fish and cavefish and in both light and dark conditions, where fish may use both auditory and tactile/lateral line sensing (in the dark) in addition to visual sensing (in the light in surface fish). However, chemical stimuli (food extract) evoked a prominent foraging response in both surface fish and cavefish for both light and dark conditions than the object stimuli (beads). In the dark, both morphs directly aimed at the bottom of the tank (food extract does not stimulate visual sensation), where their food always ended up, suggesting chemical stimuli did not navigate them toward food sources but instead evoked fish to the existence of food. Cavefish showed higher foraging activities than surface fish under chemical stimulus.

In summary, surface fish were visually driven and tended to require multiple sensory stimuli to evoke foraging. In contrast, the sole auditory stimulus was still able to evoke foraging behavior in cavefish. Among the given stimuli, chemical stimulus strongly drove foraging behavior immediately at the bottom of the tank and/or at the water surface in both surface fish and cavefish whilst the food extract plume was still at the middle of the water column, suggesting fish did not directly use chemical gradients but instead used this stimulus as ambient cues and searched where food was likely to exist. Further, we also detected different foraging patterns between the light and dark conditions even in blind cavefish, and the differences in diet-locating strategies—zigzag and circling—between surface fish and cavefish. Our result provides new evolutionary insight into foraging strategies for diet-related stimuli.

## Materials and Methods

### Fish maintenance and care

Populations of *A. mexicanus* (both sighted and the blind morphs) were raised and bred at the University of Hawai’i at Mānoa aquatic facility with care and protocols approved under IACUC (17-2560) at University of Hawai’i at Mānoa. Both surface fish and cavefish were *Astyanax mexicanus* species. Surface fish raised in the lab were descendants from those collected by Dr. William R. Jeffery from Balmorhea Springs State Park in Texas and cavefish were descendants collected by Richard Borowsky and Dr. William R. Jeffery in Cueva de El Pachón in Tamaulipas, Mexico. Both surface fish and cavefish were raised on a 12:12 light cycle in 42-liter tanks in a custom water-flow tank system. Temperatures were maintained at 21ºC ± 0.5ºC for rearing, 24ºC ± 0.5ºC for behavior experiments, and 25ºC ± 0.5ºC for breeding. Their diet consisted of TetraColor tropical fish food granules and TetraMin tropical fish food crisps, tetra, Blacksburg, VA, and jumbo mysis shrimp (Hikari Sales, USA, Inc., Hayward, CA). Fish were fed on Zeitgeber time 3 and 9 and maintained at 7.0 pH with a water conductivity of 600–800 μS.

### Experimental populations

We used a 37.9 L tank to house each experimental population (surface and cavefish) prior to introducing the stimuli. Four days prior to recording, fish tanks were cleaned and the tank water was replaced with conditioned fish water (pH 6.8–7.0, conductivity: ∼700 μS adjusted with Reef Crystals Reef Salt, Instant Ocean, Blacksburg, VA). At least three days prior to recording, fish circadian rhythm was entrained by a 12:12 h light-dark cycle with 30–100 lux light. On recording days, the experiment commenced at ∼2 hours of Zeitgeber time. We used a 10-min acclimation time prior to recording. Each 37.9 L tank contained three replicate fish (N = 3). The stimuli were administered in the following order: (1) water droplets (3 drops), (2) red plastic beads (4.7 mm diameter: Millipore Sigma, Burlington, MA), (3) food extract (see below), (4) a combination of food extract & beads, and (5) agar-solidified food (see below). Each of the stimuli were given in 10-min intervals. Recording was performed for ∼50 min in total. The dark experiment (no light) and the light experiment (30–100 lux) were performed on different days.

### Experimental stimulus

The water stimulus was three droplets of distilled water and 4–5 of red polystyrene beads (4.7mm in diameter). The food extract was made by suspending 0.1 g of fine ground Tropical XL Color Granules with Natural Color Enhancer (Tetra U.S., Blacksburg, VA) in 2 mL of distilled water mixed with 0.5 mL of 0.5% Methylene Blue (MilliporeSigma) and filtered with a 0.45 μm syringe filter. The food extract was made fresh for each experiment and three drops were added as the stimulus. The agar-solidified food was comprised of 1.0 g of fine ground Tropical XL Color Granules with Natural Color Enhancer (red colored granules) suspended with 5 mL of 1% agar (MilliporeSigma) in the fish conditioned water (pH 6.8–7.0, conductivity ∼700 μS), then poured into 6-cm dishes to solidify. Once solidified, a razor blade sterilized with 70% ethanol was used to cut the agar food into 5 × 5 mm squares and 3–4 pieces were given per stimulus. Sinking of red plastic beads was approximately the same as the red agar food, mimicking red agar food movement.

### Recordings

All light condition videos were recorded on an iPhone Xs (Apple, Cupertino, CA) at 30 fps. Fish behaviors in the dark were recorded using a custom-made infrared back-light system (SMD 3528 850nm strip: LightingWill, Guang Dong, China). A LifeCam studio 1080p HD webcam (Microsoft, Redmond, WA, USA) with a zoom lens (Zoom 7000, Navitar, Rochester, NY, USA) fitted with an IR high-pass filter (Optical cast plastic IR long-pass filter, Edmund Optics Worldwide, Barrington, NJ, USA). A USB webcam (LifeCam studio 1080p HD webcam, Microsoft, Redmond WA, US) was used to record at 16–20 fps using virtual dub software (version 1.10.4, http://www.virtualdub.org/). Once recorded, videos were uploaded to Google Drive for accessibility.

### Video analysis

Videos were analyzed using Behavioral Observation Research Interactive Software (BORIS V. 7.4.11-2019-02-28, Department of Life Sciences & Systems Biology, University of Torino-Italy). For video analysis, the tank was divided into nine square sections, with areas 1, 2, 3, and 5 as the top row and areas 7–9 as the bottom (Fig 1B, the far-left panel). Using BORIS, each fish’s actions were recorded during the videos. Latency was defined as the measurement of time duration between when stimulus hit the water surface and when fish of interest approached at the dropping point. “Attempts” were measured as the number of capturing or biting motion against the stimulus by observing the opening and closing of the mouth rapidly or picking up a bead/food. A “zigzag” motion was defined as rapid changes of the swimming direction every ∼ 1 s and was measured as occurrence (bout number) and duration (s). “Circling” motion was defined as the continuous unidirectional turnings without glide swimming, and was measured as occurrence (bout number) and duration (s) by unidirectional turning to make at least one full circle at the tank bottom or water surface.

**Fig 1.**
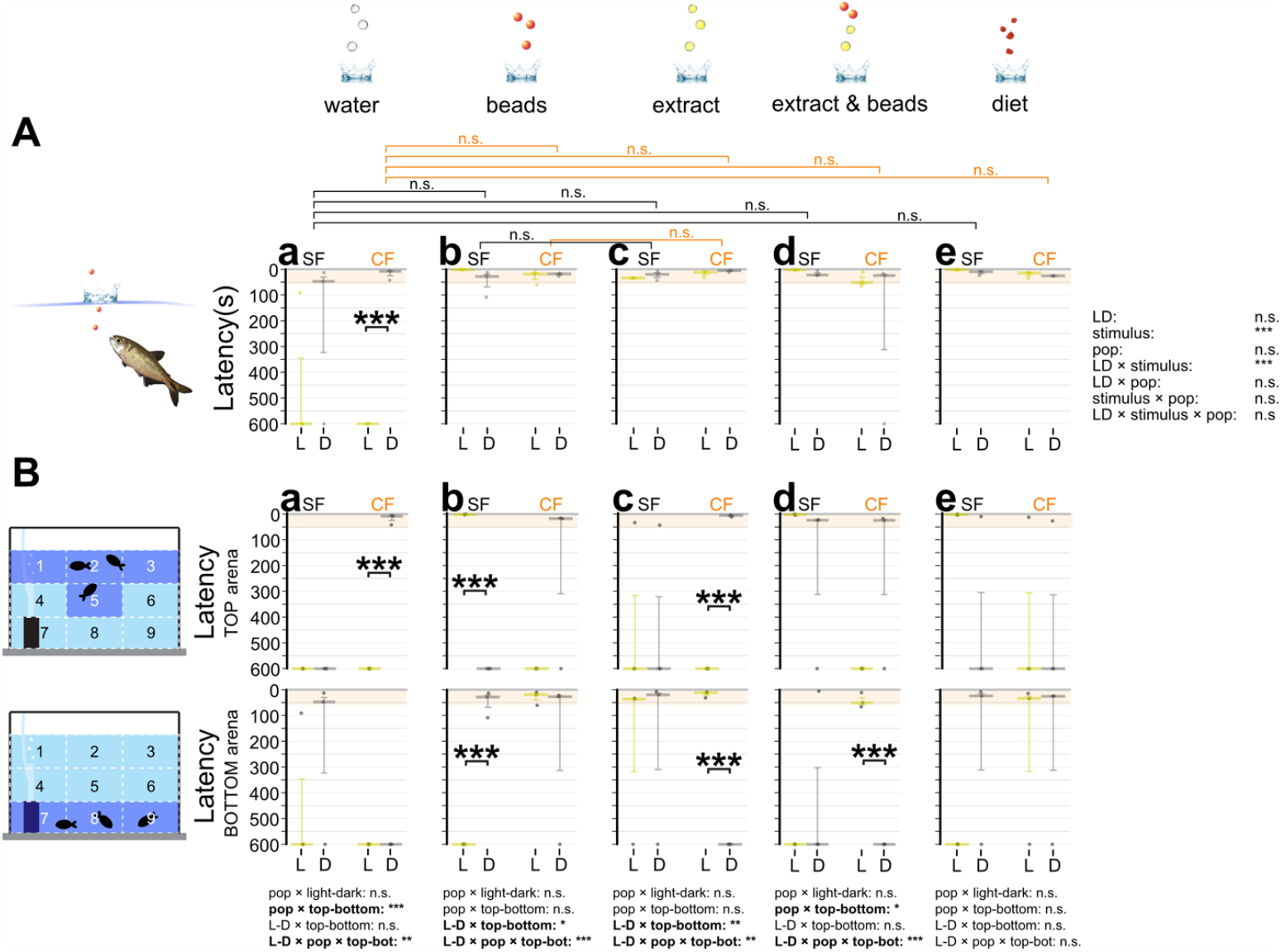
Latencies in response times to different sensory stimuli. (**A**) Overall latency (s) between when the object hit the water surface and when fish directly aimed toward the object. Three fish in a tank were given three droplets of reverse-osmosis (RO) purified water (water: panels **Aa** and **Ba**), three red plastic beads 4.7 mm in diameter (beads: **Ab** and **Bb**), three droplets of food extract (extract: **Ac** and **Bc**), three droplets of food extract followed by three red beads (extract and beads: **Ad** and **Bd**), and 3–4 granules (3–5 mm in diameter) of actual food (diet: **Ae** and **Be**; see Materials and Methods). (**A**) Latencies of surface fish (SF: left) and cavefish (CF: right) are shown on the y-axis. Top: shorter latency; bottom: no response within a 10 min observation (600 s). Latencies under light conditions (L: yellow bars and dots) and dark conditions (D: gray bars and dots) are also shown. The first 60 seconds after the object hit the water surface are shaded red. The statistical test results of the generalized linear model are shown on the far right. For each comparison, light and dark conditions were compared within the population per treatment (e.g., a bracket in CF with the water stimulus). Within each population, different stimuli were compared with the water stimulus and significances were calculated via Mann-Whitney tests adjusted by Holm’s correction, shown as brackets at the top of boxes. All comparisons were non-significant (n.s.) in latencies. (**B**) Fish locations were tracked as the top (top row) or bottom (bottom row) and measured latencies. The far-left panels indicate the areas counted as the top (areas 1, 2, 3 and 5), and the bottom (areas 7, 8 and 9). The y-axes and brackets in **Ba**-**Be** represent the same as (**A**). All stars represent P-values after Holm’s correction. Statistical test summaries using the generalized linear model including arena locations (top-bottom) are shown at the bottom of the boxes. Only interaction results are shown. Details of all statistics scores in this figure are found in Supplementary Data 1. n.s.: not significant, *: P < 0.05, **: P < 0.01, ***: P < 0.001.

We recorded the tank areas where each behavior was observed. Quantitative data collected from BORIS was then consolidated in the Excel macro (Microsoft, Redmond, WA) (https://zenodo.org/record/7996590).

### Statistical analysis

Quantitative data were exported from BORIS to Excel. Using macros in Excel, data were compiled and the totals of each foraging behavior were calculated (shared on Zenodo: https://zenodo.org/record/7996590). All statistical analyses were performed in RStudio 4.0.3 (RStudio, Boston, MA, USA). The R packages used included *lme4, lmerTest, car, coin, yarrr, ggplot2, AICcmodavg*, and *ggpubr*. Linear or generalized linear models were selected using Akaike’s information criterion function to identify the best fit models for analyses for latency, attempt, and zigzag and circling motions. We used multifactorial variance analyses using generalized linear model fitting functions (glm or glmer in the *lme4* package). Post-hoc tests were performed using the Wilcoxon signed-rank test followed by Holm’s multiple-test correction.

## Results and Discussion

Foraging attempt was composed of initial investigation (measured by latency), adherence to the stimulus source (proxy of the number of attempts) and searching mode (zigzag or circling motion) to analyze differences in sensory modality between surface fish and cavefish.

### Latency

For the response to the water droplet stimulus, there was no detectable difference between surface fish and cavefish, yet we detected different responses between light and dark conditions in cavefish (water droplets; Fig 1Aa and Supplementary File 1). Detailed scoring further revealed that cavefish were attracted to water droplet stimulus when droplets hit the water surface (top) in the dark (Fig 1Ba). In contrast, under light conditions, cavefish did not respond to the water droplet. Since cavefish seem to sense ambient light with brain opsins [13] and light conditions pose increased exposure risk to the surrounding environment [14], cavefish may have a reserved response under light conditions. Surface fish did not respond to water droplets, suggesting auditory stimulus was not sufficient to evoke foraging behaviors.

For beads, which potentially stimulate visual, auditory (when it hit water surface), and tactile (when fish touched it at the bottom) sensors, surface fish responded quickly (∼10 s) by swimming toward the top and toward the bottom of the arena under light and dark conditions, respectively (Fig 1Ab and 1Bb). The latter result indicates that surface fish responded to beads without visual stimulus. This response in the light seems primarily driven by visual stimulus. In contrast, these initial responses in the dark suggest surface fish used auditory (at the top of the arena), lateral line and/or tactile sensing (at the bottom) to locate stimulus sources in the dark (Fig 1Bb). Cavefish responded to beads similarly to surface fish in the dark irrespective of light or dark conditions (Fig 1Bb), suggesting surface fish and cavefish used similar sensory modalities in initial responses against solid food-like objects in the dark.

Using food extract showed somewhat similar results to water droplets but showed strong engagement toward the bottom (surface fish in the light and dark and cavefish in the light) or the top (cavefish in the dark) (Fig 1Ac and 1Bc). Importantly, food extract always dispersed in the middle of the recording tank and the dense food-extract plume (dye with methylene blue; see Materials and Methods; Movie 1) never reached the bottom before dispersing, suggesting chemical stimulus was not used to orient food location, but may be used as a signal of food existence in a given environment (ambient existence). Cavefish aimed at the top of the tank in the dark could be explained similarly to that evoked by water droplets (i.e., boldness in the dark; see above), but significantly responded and aimed to the bottom in the lighted condition, which was not observed with the water droplet stimulus (Fig 1Bc).

The combined bead and food-extract stimulus invoked the summed response of beads-only and food extract-only stimulus in cavefish, which responded to the stimulus by either aiming to the bottom (light) or top (dark; Fig 1Bd). Surface fish were engaged toward the top under light conditions and aimed at either the top or bottom under dark conditions, which was also similar to food stimulus (Fig 1Bd and 1Be). Cavefish aimed at either the top or bottom with food stimulus and no notable difference in the feeding was detected compared with the food extract (Fig 1Bc and 1Bd).

In summary, water droplet stimulus (auditory) evoked a light-dependent response in the blind cavefish, whereby dark conditions seemed to make cavefish bold to come to the water surface. Other stimuli induced different light- and area-dependent responses in surface fish and cavefish, but opposite responses: surface fish foraged in the light, but cavefish foraged in the dark, assuming attraction to the top area as a bolder response. However, overall latencies were similar between surface fish and cavefish in different stimuli and under dark conditions (Fig 1A), suggesting cavefish did not evolve particular sensory responses during initial foraging attempts in the dark.

### Number of foraging attempts

Fish attempted to bite or capture the stimulus source following initial contact. We measured this engagement to foraging defined by darting/thrusting and biting motions against the stimulus source (i.e., attempts). In contrast to the initial response (i.e., latency), water droplets did not evoke any attempts in either surface fish or cavefish in either light or dark conditions (Fig 2Aa). All other stimuli led to significantly more attempts in both surface fish and cavefish (Fig 2Ab-Ae). For the bead stimulus, as expected, surface fish were well engaged by showing more attempt numbers than water droplets under light conditions (both at the top and bottom of the recording arena; Fig 2Bb), but still responded to dark conditions (at the arena bottom; Fig 2Ab and 2Bb). Surface fish responses in the dark may be based on tactile or lateral line sensors since surface fish attempted to bite beads only close to or when touching beads (1–2 cm), which is the sensing range of tactile and lateral line sensors. Chemical sensing is likely not involved here because beads did not emit food-like chemicals. Most surface fish mouthed beads, suggesting chemical stimulus—typically detected by extra mouth taste buds [15,16]—is not necessary involved in capturing ‘food’-like objects. Cavefish were less attracted to beads (effect size, r = 0.66 compared with surface fish’s r = 0.82; Fig 2Ab), but showed more attempts compared with water droplets (Fig 2Ab). Some cavefish showed a number of attempts at the top tank area in the dark (Fig 2Bb). Cavefish attempts in the top tank area could be based on similar reasons as latency: using auditory input and being bold in the dark. Cavefish did not show many attempts for beads in the bottom tank area under light or dark conditions compared with surface fish (Fig 2Bb), suggesting cavefish may need additional stimuli, such as chemicals. In summary, cavefish may need further sensory inputs (integrating alternative sensory inputs) in addition to the object stimulus to maintain foraging behavior compared with surface fish.

**Fig 2.**
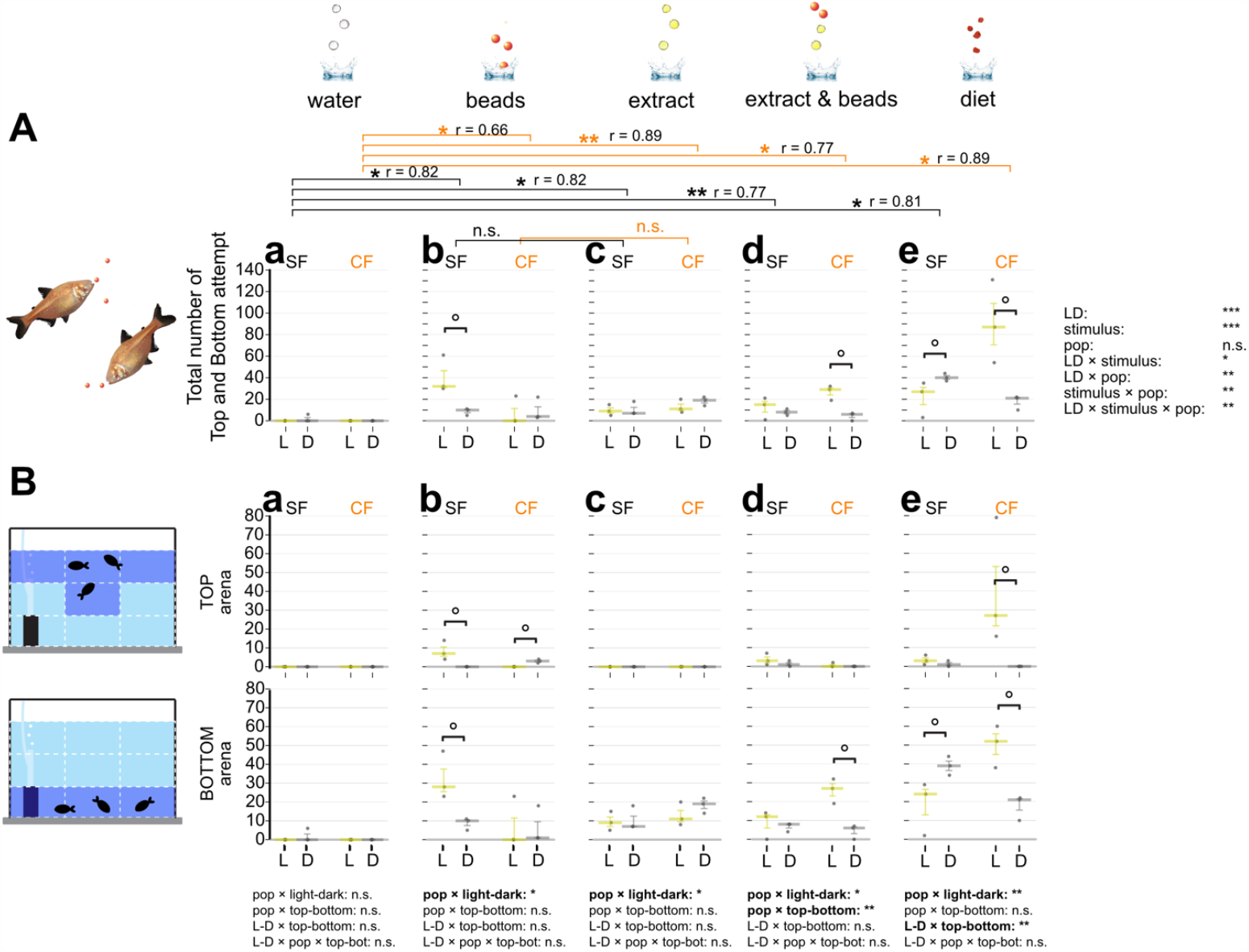
Measured attempts responding to different sensory stimuli. Overall attempt number in the 10-minute experiment defined as when fish obviously attempted a strike at the stimulus within the top or bottom areas. Three fish in a tank were given three droplets of RO purified water (water; **Aa** and **Ba**), three red plastic beads 4.7 mm in diameter (beads; **Ab** and **Bb**), three droplets of food extract (extract; **Ac** and **Bc**), three droplets of food extract followed by three red beads (extract and beads; **Ad** and **Bd**), and 3–4 granules (3–5 mm in diameter) of actual diet (diet; **Ae** and **Be**) (see Materials and Methods). In **Aa**-**Ae**, attempt(s) of surface fish (SF: left) and cavefish (CF: right) are plotted on the y-axis. Attempts under light condition (L: yellow bars and dots) and dark condition (D: gray bars and dots) are also shown. Statistical test result of the generalized linear model are shown on the far right (**A**). For each comparison, light and dark conditions were compared within the population per treatment as in Fig 1. Within each population, different stimuli were compared with the water stimulus and significances were calculated via Mann-Whitney tests adjusted by Holm’s correction, shown as brackets at the top of the boxes. Comparisons between light and dark and between stimuli were significant. We also found significant differences when comparing light and dark responses and the stimuli and several interactions among the stimuli, populations, and light conditions. Details are available in Supplemental Table 1. (**B**) Fish locations were tracked as the top (top row) or bottom (bottom row) and measured attempts. The Y-axes and brackets represent the same as (**A**). All stars represent P-values after Holm’s correction. Statistical test summaries using the generalized linear model including arena locations (top-bottom) are shown at the bottom of the boxes. Only interaction results are shown. Details of all statistics scores in this figure are in Supplementary Data 1. n.s.: not significant, °: P < 0.10, *: P < 0.05, **: P < 0.01, ***: P < 0.001.

Diet-extract chemical stimulus facilitated more attempts in both surface fish and cavefish irrespective of light or dark conditions (Fig 2Ac and Bc). These foraging attempts were mainly observed in the bottom tank area where food always sunk, suggesting fish may forage based on their previous experiences where the food always ended up.

For combined beads and food-extract stimulus, surface fish foraging patterns were similar to those observed in bead-only trials (see above; Fig 2Ad and Bd). However, cavefish increased their foraging attempts under light conditions probably based on higher activity under ambient light [13]. Compared with bead- and food extract-only trials, the combined stimulus with light may simultaneously facilitate foraging attempts where cavefish showed higher activities under light. This notion was supported by food stimulus where cavefish also showed high attempts under light (Fig 2Ae and 2Be). For food stimulus, cavefish were more active under light than dark conditions, which seems to contradict the result of latency measurements (Fig 2Ae and 2Be; Fig 1Ba-d). However, these and the latency results indicate cavefish could be more alert with light during initial approaches, but higher cavefish activity under light could have resulted in more attempts toward the stimulus source.

Surface fish showed higher attempts in dark than light conditions (Fig 2Ae and 2Be). However, the mechanism remains unclear. One possible explanation in the food stimulus trial is that the foraging sound of their cohorts evokes foraging behaviors in others [17]. Surface fish may respond to such sounds in the dark [18,19], although cavefish may have reached at the plateau of their response to external foraging sounds. This prediction requires further testing.

### Food discovery strategy (zigzag and circling motions)

Surface fish and cavefish showed specific movement patterns to locate stimulus (food), namely zigzag and circling motions (see Materials and Methods-Video Analysis). Both patterns were observed in surface fish and cavefish but used to varying degrees and in different contexts.

#### Zigzag motion

The zigzag motion was detected mainly with chemical stimulus (food extract, combined and diet stimulus) and evoked in the dark (Figs 3 and 4). This trend changed when cavefish confronted multiple stimulus (i.e., combined beads and food extract), where cavefish showed higher instances of zigzag motion under light conditions, as well as for surface fish toward foraging sounds (agar food stimulus). In summary, this zigzag motion is a shared response in surface and cavefish primarily without visual inputs.

**Fig 3.**
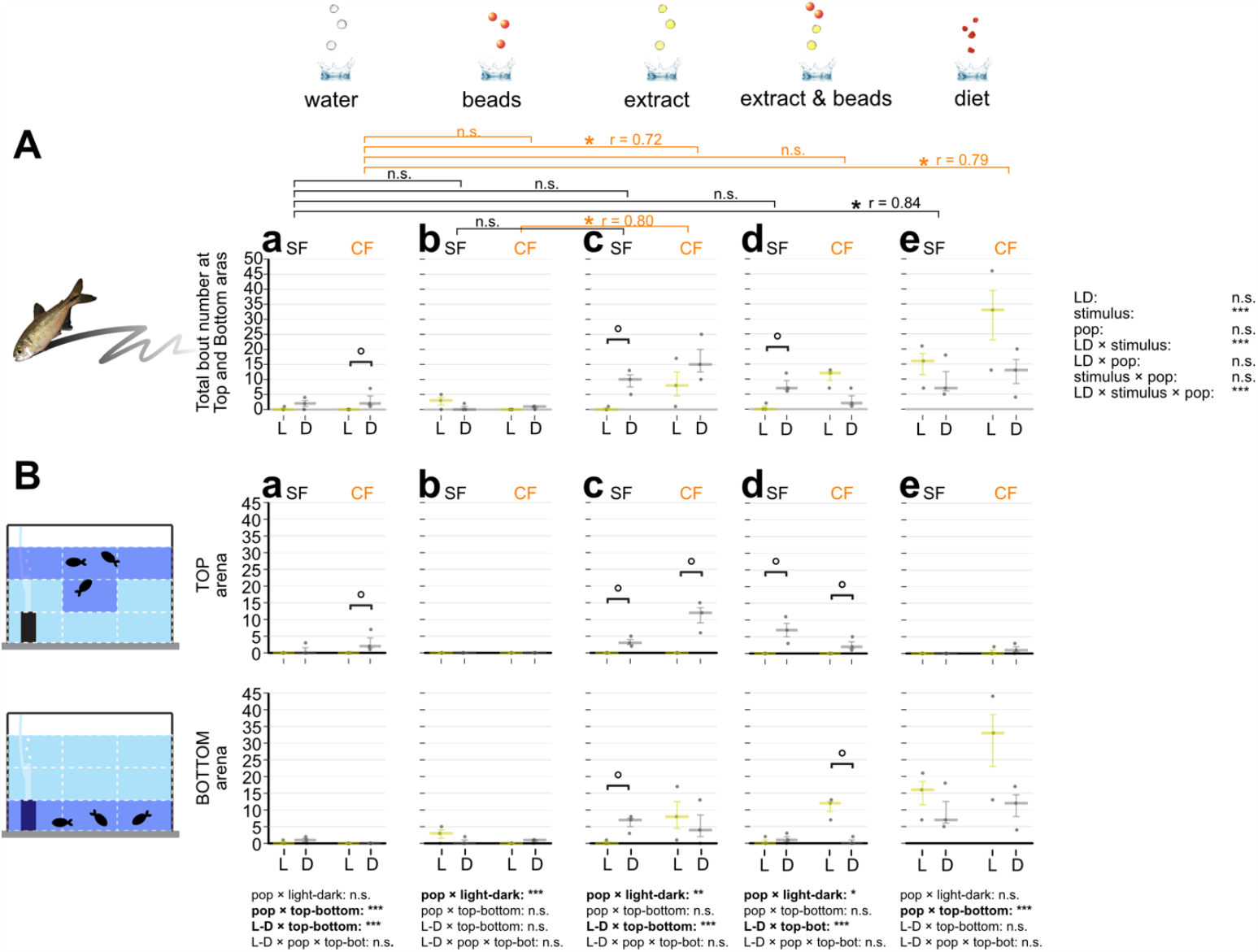
Bout number of zigzag searching behavior in response to different sensory stimuli. (**A**) Overall bout (i.e., event) counts for searching behavior using zigzag(s) in the 10-minute experiment. Zigzag searching behavior was defined as fish searching by zigzag motion (back and forth) frequently at the water surface or tank bottom with sensory stimuli (see Materials and Methods). The zigzag bout numbers of surface fish (SF: left) and cavefish (CF: right) are plotted on the y-axis. Zigzag behavior under light condition (L: yellow bars and dots) and dark condition (D: gray bars and dots) are also shown. Statistical test result of the generalized linear model is shown on the far right. For each comparison, light and dark conditions were compared within the population per treatment. Within each population, different stimuli were compared with water stimulus and significances were calculated via Mann-Whitney tests adjusted by Holm’s correction (See Supplementary Data 1). (**B**) Fish locations were tracked as the top (top row) or bottom (bottom row) and measured zigzag behavior. The y-axes and brackets represent the same as (**A**). All stars represent P-values after Holm’s correction. Statistical test summaries using the generalized linear model including arena locations (top-bottom) are shown at the bottom of the boxes. Only interaction results are shown. Details of all statistics scores in this figure are in Supplementary Data 1. n.s.: not significant, °: P < 0.10, *: P < 0.05, **: P < 0.01, ***: P < 0.001.

**Fig 4.**
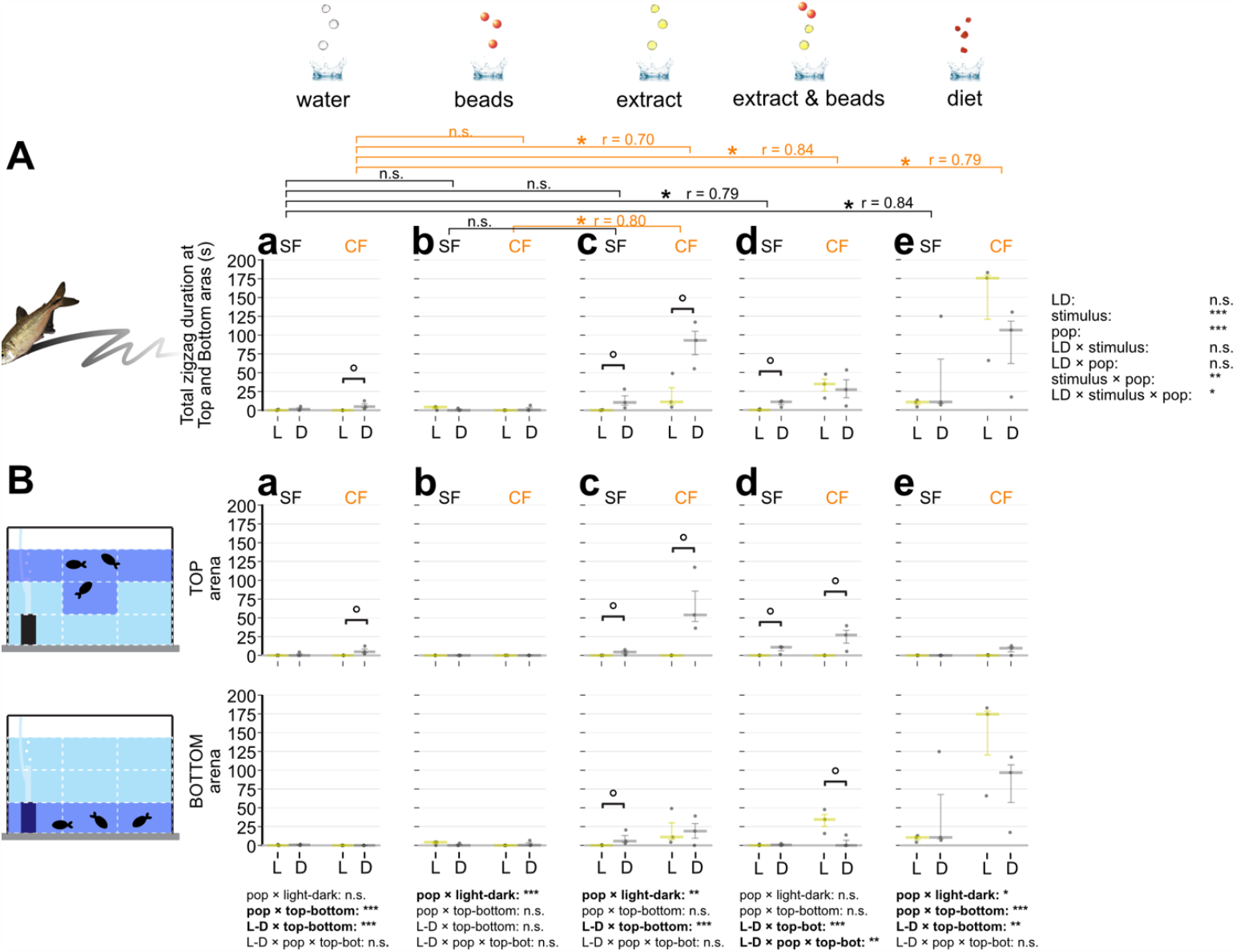
Zigzag searching duration in response to different sensory stimuli. (**A**) Overall searching duration (s) using zigzag(s) in the 10-minute experiment. Zigzag searching duration was measured when fish were searching with back-and-forth movements. The experimental setup was the same as Figs 1 and 3 (see Materials and Methods). The measured duration (s) of zigzag behavior of surface fish (SF: left) and cavefish (CF: right) are plotted on the y-axis in each panels (**Aa**-**Ae**). Zigzag behavior under light condition (L: yellow bars and dots) and dark condition (D: gray bars and dots) are also shown. Statistical test result of the generalized linear model is shown on the far right. For each panel, light and dark conditions were compared within the population per treatment. Within each population, different stimuli were compared with the water stimulus, and significances were calculated via Mann-Whitney tests adjusted by Holm’s correction (See Supplemental Data 1). (**B**) Fish locations were tracked as the top (top row) or bottom (bottom row) and measured the zigzag behavior duration. The y-axes and brackets represent the same as (**A**). All stars represent P-values after Holm’s correction. Statistical test summaries using the generalized linear model including arena locations (top-bottom) are shown at the bottom of the boxes. Only interaction results are shown. Details of all statistics scores in this figure are in Supplementary Data 1. n.s.: not significant, °: P < 0.10, *: P < 0.05, **: P < 0.01, ***: P < 0.001.

#### Circling motion

The circling motion was observed mainly with chemical stimulus as seen in the zigzag motion, but was more dominant in cavefish than surface fish (Figs 5 and 6). Cavefish exhibited high levels of circling motion under light conditions with chemical stimuli (food extract and combined beads and food extract). Circling could be a better strategy than zigzagging given that circling yields fish come nearby the same food multiple times while only once while zigzagging.

**Fig 5.**
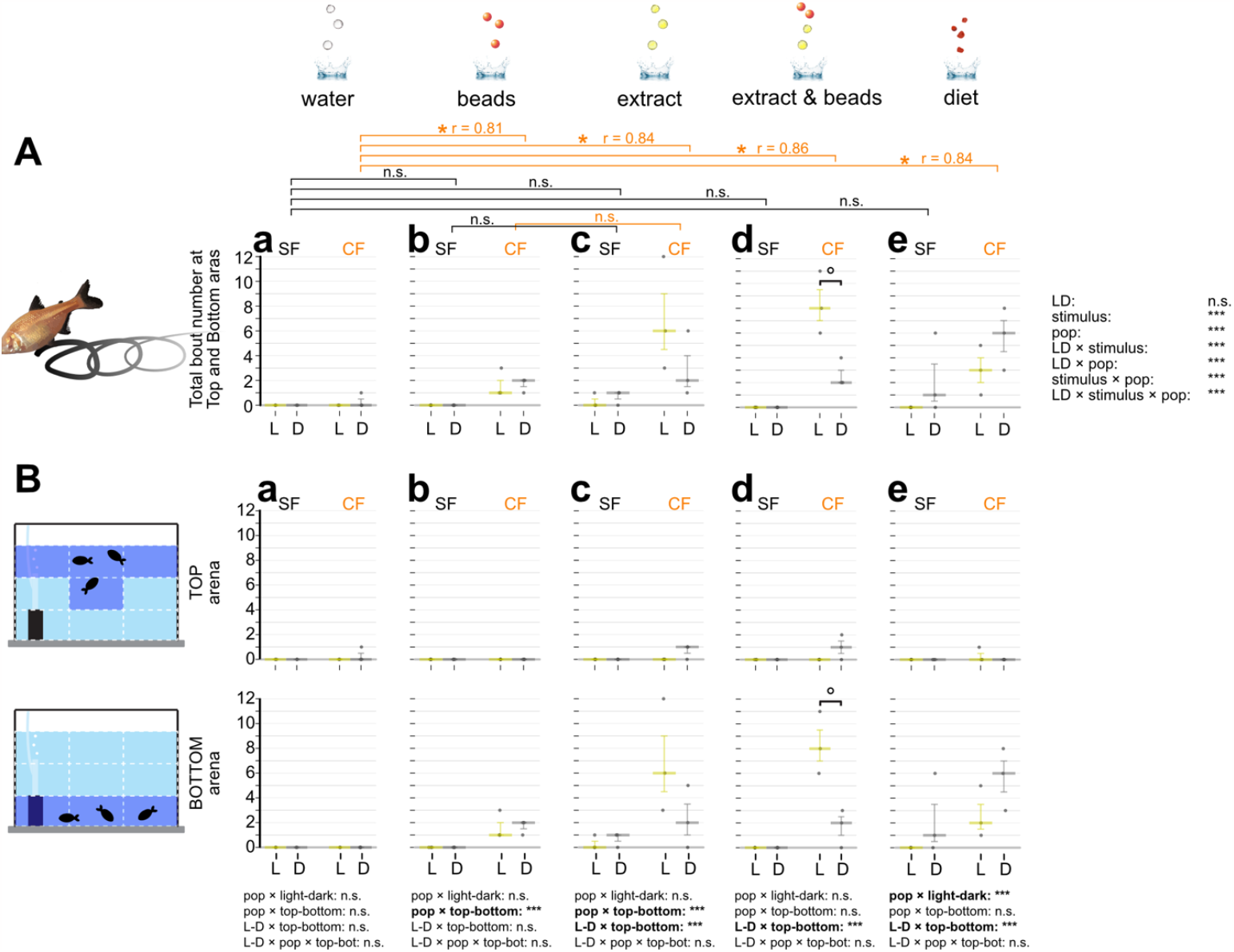
Bout numbers of circling searching behavior in response to different sensory stimuli. (**A**) Overall bout (i.e., event) numbers of circling motions fish during the 10-minute assay. Circling searching behavior is defined as fish repeating a circle pattern. The stimuli were given as in Fig 1 (see Materials and Methods too). The bout numbers of the circling motions of surface fish (SF: left) and cavefish (CF: right) were plotted on the y-axis during a 10-min observation in each panel of **Aa**-**Ae**. Circling behavior under light condition (L: yellow bars and dots) and dark condition (D: gray bars and dots) are also shown. Statistical test result of the generalized linear model is shown on the far right. For each comparison, the light and dark conditions were compared within the population per treatment. Within each population, different stimuli were compared with the water stimulus and significances were calculated via Mann-Whitney tests adjusted by Holm’s correction (see Supplementary Data 1 too). (**B**) Fish locations were tracked as the top (top row) or bottom (bottom row) and measured circling behavior. The y-axes and brackets represent the same as (**A**). All stars represent P-values after Holm’s correction. Statistical test summaries using the generalized linear model including arena locations (top-bottom) are shown at the bottom of the boxes. Only interaction results are shown. Details of all statistics scores in this figure are in Supplementary Data 1. n.s.: not significant, °: P < 0.10, *: P < 0.05, ***: P < 0.001.

**Fig 6.**
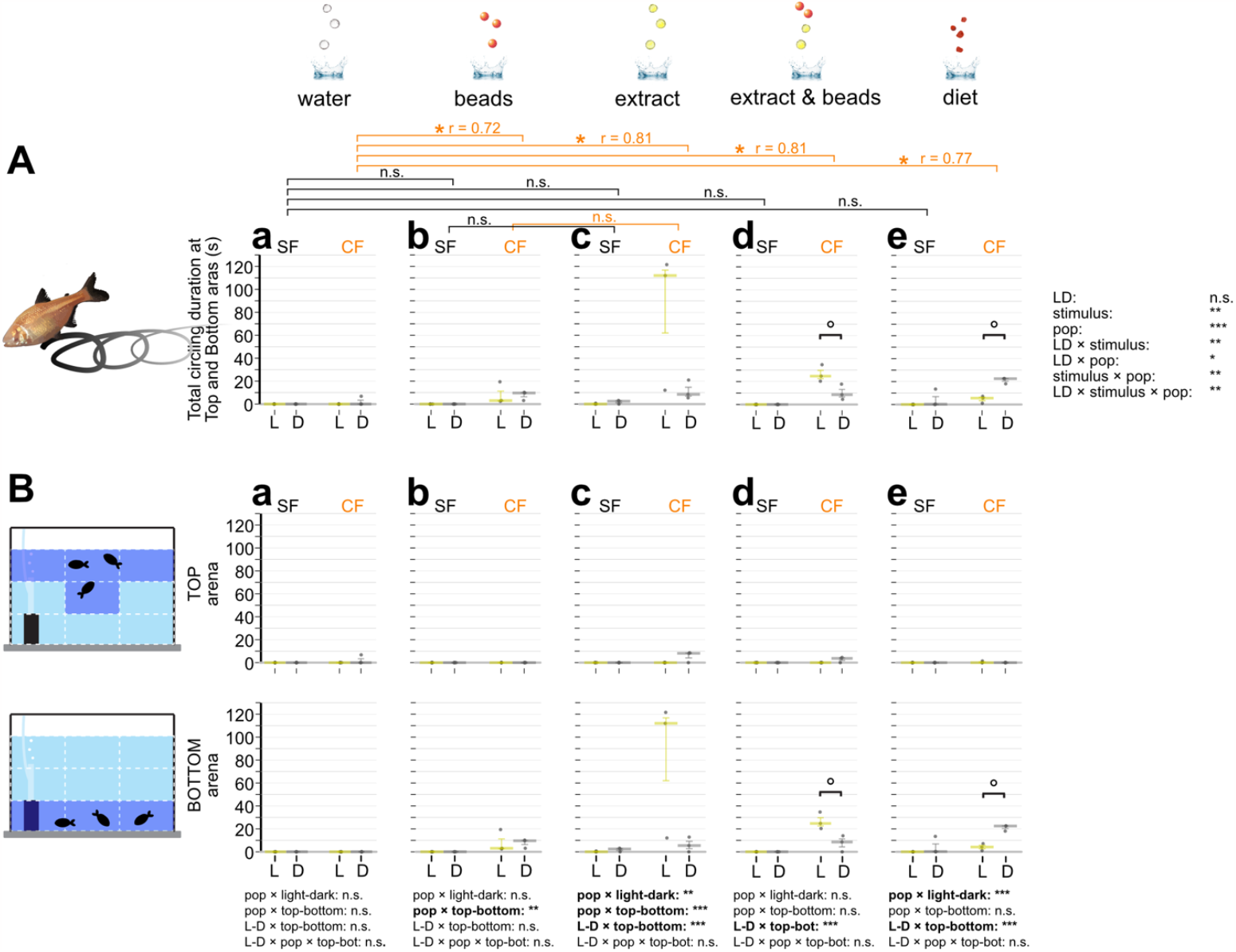
Circling searching duration in response to different sensory stimuli. (**A**) Overall duration of searching showing circling during the 10-minute observation. Circling searching duration is defined from when fish began searching in a repeated circle pattern to when fish stopped the behavior. Stimuli were given as in Figs 1 and 5 (see Materials and Methods). Duration of circling behavior of surface fish (SF: left) and cavefish (CF: right) were plotted on the y-axis within a 10 min observation in each panel of **Aa**-**Ae**. Circling behavior under light condition (L: yellow bars and dots) and dark condition (D: gray bars and dots) are also shown. Statistical test results of the generalized linear model are shown on the far right. For each panel, light and dark conditions were compared within the population per treatment. Within each population, different stimuli were compared with the water stimulus and significances were calculated via Mann-Whitney tests adjusted by Holm’s correction, shown as brackets at the top of the boxes (see also Supplemental Data 1). (**B**) Fish locations were tracked as the top (top row) or bottom (bottom row) and measured circling behavior time. The y-axes and brackets represent the same as (**A**). All stars represent P-values after Holm’s correction. Statistical test summaries using the generalized linear model including arena locations (top-bottom) are shown at the bottom of the boxes. Only interaction results are shown. Details of all statistics scores in this figure are in Supplementary Data 1. n.s.: not significant, °: P < 0.10, *: P < 0.05, **: P < 0.01, ***: P < 0.001.

## Conclusion

We examined foraging responses of surface and cavefish using water droplets (auditory stimulus), plastic beads (visual+auditory+lateral line+tactile), food extract (auditory+chemical), plastic beads & food extract, and actual food. To maximize foraging efficiency and minimize energy loss, visual/light conditions for surface fish favored beads and actual food (low latency; Fig 1) and surface fish captured these sources with a low number of attempts (Fig 2Ab, 2Ad, 2Ae, 2Bb, 2Bd and 2Be). Surface fish could also conserve energy by reducing total attempts toward non-visible objects (water droplets; Fig 2Aa and 2Ba). In contrast, in the dark, both surface and cavefish responded to auditory stimulus (water droplets; Fig 1Aa and 1Ba) to investigate without performing extra attempts (fewer attempts in water droplets; Fig 2Aa and 2Ba), which may be an efficient strategy to investigate objects if it is food. However, surface fish were less efficient with plastic beads by showing much higher attempts toward this inedible object (Fig 2Ab and 2Bb) than cavefish, suggesting visual stimulus is highly favored in foraging. In contrast, chemical stimulus evoked a higher number of attempts in cavefish than surface fish, indicating higher sensory emphasis on chemical sensing (olfaction and taste buds) for foraging in cavefish. This sensory priority in olfaction in cavefish is supported by the previous report indicating that cavefish responded to 10^5^ times lower concentrations of amino acid stimulus (10^−5^ M vs 10^−10^ M of alanine in surface fish vs cavefish, respectively [10]. However, neither cavefish nor surface fish appeared to use chemical stimulus to navigate themselves toward sources as cavefish (and surface fish in the dark) started searching for food at the water surface or tank bottom immediately after touching food extract clouds in the middle of the water column (Movie 1), suggesting chemical stimulus indicated food presence instead of that fish use the odor gradient. This feeding strategy seems to contradict the previous reports where the chemical gradient looked to navigate *Astyanax* fish [10,20]. However, we suspect that, while the chemical gradient informs the approximate direction that the fish must swim to approach the source of food in a still-water pool [20], the precise location of any suspended food particle is difficult to identify based on chemical sensing because of the slow diffusion of molecules, which are advected by the fluid flow over a long time before they reach the fish’s chemoreceptors. In contrast, the relatively fast diffusion of momentum through the viscous boundary layer around the fish enables particles near the boundary layer to be located quickly based on mechanical sensing [21]. Further study is needed to confirm this in a noisy environment.

Cavefish were more active by showing more attempts under light than dark when food scent was available (food extract and agar food), possibly due to higher activity under light [13] while foraging behavior was evoked by chemical stimulus (Fig 2Ad and 2Ae). We suspect this light-dependent response in cavefish is due to an evolutionary artifact of ambient light detection based on non-ocular opsins [13].

While both surface fish and cavefish showed similar levels of zigzag foraging in the dark (Figs 3 and 4), cavefish exhibited much more circling foraging than surface fish (Figs 5 and 6), suggesting circling may be an evolutionarily-enhanced strategy in cavefish, i.e. food could be less dispersed at the tank bottom compared with zigzagging, and also, cavefish have more chances to sense the same food multiple times compared with zigzagging, yielding only once in given time. This idea needs further investigation to measure differences in foraging efficiency between zigzagging and circling.

## Acknowledgments

We are grateful to V Crystal, J Choi, L Lu, J Nguyen, C Balaan, K Lactaoen, M Worsham, H Hernandez, N Doeden, J Kato, M Ito, R Balmilero-Unciano, E Doy, A Martinez, D Mones, H Yoshizawa for fish care assistance. We also thank to M Iwashita for reviewing the final version of manuscript. We gratefully acknowledge support from the National Institute of Health (P20GM125508) to MY, Hawaii Community Foundation (18CON-90818) to MY.

## Author contributions

KK: designed the experiments, performed the experiment and analyses, wrote the initial draft, and edited the manuscript.

VFLF: designed the experiments, performed and assisted the experiment, and edited the manuscript.

DT: designed the experiment, consulted the experiment, and edited the manuscript.

MY: designed the experiments, performed the experiment and analyses, wrote the initial draft with KK, and edited the manuscript

## Data Availability

The video datasets generated and/or analyzed during the current study are available at the university’s shared server and will be deposited to Zenodo (https://zenodo.org/). The MS Excel macro used in this study is available at https://zenodo.org/record/7996590

